# Constant Infusion Radiotracer Administration for High Temporal Resolution Positron Emission Tomography (PET) of the Human Brain: Application to [18F]-Fluorodexoyglucose PET (FDG-PET)

**DOI:** 10.1101/667352

**Authors:** Sharna D Jamadar, Phillip GD Ward, Alexandra Carey, Richard McIntyre, Linden Parkes, Disha Sasan, John Fallon, Shenpeng Li, Zhaolin Chen, Gary F Egan

## Abstract

Functional Positron Emission Tomography (fPET) provides a method to track molecular dynamics in the human brain. With a radioactively labelled glucose-analogue, [18F]-flurodeoxyglucose (FDG-fPET), it is now possible to index the dynamics of glucose metabolism with temporal resolutions approaching those of functional magnetic resonance imaging (fMRI). This direct measure of glucose uptake has enormous potential for understanding normal and abnormal brain function, and probing the effects of metabolic and neurodegenerative diseases. Further, new advances in hybrid MR-PET hardware makes it possible to capture fluctuations in glucose and blood oxygenation simultaneously using fMRI and FDG-fPET.

The temporal resolution and signal-to-noise of the FDG-fPET images is critically dependent upon the administration of the radioactive tracer. In this work we present two alternative continuous infusion protocols and compare them to a traditional bolus approach. We detail a method for acquiring blood samples, time-locking PET, MRI and experimental stimulus, and administrating the non-traditional tracer delivery. By applying a visual stimulus, we demonstrate cortical maps of the glucose-response to external stimuli on an individual level with a temporal resolution of 16-seconds.

**Summary:** Radiotracer infusion protocols for positron emission tomography (PET) provide improved temporal resolution over bolus administration. Here, we describe radiotracer administration for two protocols, constant infusion and bolus plus infusion protocol. We compare this to the standard bolus administration protocol. Using [18-F] fluorodeoxyglucose PET (FDG-PET) as an example, we show that temporal resolutions of approximately 16sec are achievable using these protocols.

## Introduction

Positron emission tomography (PET) is a powerful molecular imaging technique that is widely used in both clinical and research settings (see Heurling et al., 2017 for a recent comprehensive review). The molecular targets that can be imaged using PET are only limited by the availability of radiotracers, and numerous tracers have been developed to image neural metabolism, receptors, proteins and enzymes (Chen et al., 2018; Jones & Rabiner, 2012). In neuroscience, one of the most used radiotracers is [18F]-fluorodeoxyglucose (FDG-PET), which provides an index of cerebral glucose metabolism. The human brain requires a constant and reliable supply of glucose to satisfy its energy requirements (Kety, 1957; Sokoloff, 1960), and 70-80% of cerebral glucose metabolism is used by neurons during synaptic transmission (Harris, Jolivet, & Attwell, 2012). Changes to cerebral glucose metabolism are thought to initiate and contribute to numerous conditions, including psychiatric, dementia, neurodegenerative, and ischaemic conditions (Mosconi et al., 2009; Pagano, Niccolini, & Politis, 2016; Petit-Taboue, Landeau, Desson, Desgranges, & Baron, 1998). Furthermore, as FDG uptake is proportional to synaptic activity (Chugani, Phelps, & Mazziotta, 1987; Phelps & Mazziotta, 1985; Zimmer et al., 2017), it is considered a more direct and less confounded index of neuronal activity, compared to the more widely used blood oxygenation level dependent functional magnetic resonance imaging (BOLD-fMRI) response.

The majority of existing FDG-PET studies in the human brain have acquired ‘static’ images of cerebral glucose metabolism. In this method, the participant rests quietly for 10-mins with their eyes opened in a darkened room. The full dose is administered as a bolus over a period of seconds, and the participant then rests for a further 30-mins. Following the uptake period, participants are placed in the centre of the PET scanner, and a PET image is acquired, which reflects the cumulative FDG distribution over the course of the uptake and scanning periods. Thus, the neuronal activity indexed by the PET image represents the cumulative average of all cognitive activity over uptake and scan periods, and is not specific to cognitive activity during the course of the scan. This method has provided great insight into the cerebral metabolism of the brain and neuronal function. However, the temporal resolution is ~45mins, effectively yielding a static measurement in comparison to cognitive processes and common experiments in neuroimaging. Due to the limited temporal resolution, the method provides a non-specific index of glucose metabolism (i.e., not locked to a task or cognitive process) and cannot provide measures of within-subject variability (which can lead to erroneous scientific conclusions due to Simpson’s Paradox^1^; Roberts, Hach, Tippett, & Addis, 2016). Furthermore, recent attempts to apply connectivity measures to FDG-PET can only measure ‘connectivity’ across subjects; thus, differences in connectivity can only be compared between groups and cannot be calculated for individual subjects. While it is debatable *what* exactly across-subject connectivity actually measures (Horwitz, 2003), the poor temporal resolution of the bolus FDG-PET method means that ‘metabolic connectivity’ cannot be used as a biomarker for disease states, or to examine the source of individual variation, as has been previously attempted (e.g., Yakushev, Drzezga, & Habeck, 2017).

In the past 5 years, the development and wider accessibility of clinical-grade simultaneous MRI-PET scanners has sparked renewed research interest in FDG-PET imaging (Chen et al., 2018). With these developments, researchers have focused on improving the temporal resolution of FDG-PET to approach the standards of BOLD-*f*MRI (between ~0.5-2.5sec). Dynamic FDG-PET acquisitions, which are often used clinically, use the bolus administration method and reconstruct the list-mode data into bins. The bolus dynamic FDG-PET method offers a temporal resolution of around 100sec (e.g., Tomasi et al., 2017); this is clearly much improved compared to static FDG-PET imaging, but is not comparable to BOLD-fMRI. Additionally, the window in which brain function may be examined is limited as the blood plasma concentration of FDG diminishes.

To expand this experimental window, a handful of studies (e.g., Hahn et al., 2016; Hahn et al., 2018; Jamadar et al., 2019; Rischka et al., 2018; Villien et al., 2014) have adapted the radiotracer infusion method previously proposed by Carson (2000; 1993). In this method, which is sometimes described as ‘functional FDG-PET’ (FDG-*f*PET; analogous to BOLD-*f*MRI), the radiotracer is administered as a constant infusion over the course of the entire PET scan (~90mins). The goal of the infusion protocol is to maintain a constant plasma supply of FDG to track dynamic changes in glucose use across time. In a proof-of-concept study, Villien et al. (2014) used a constant infusion protocol and simultaneous MRI/FDG-*f*PET to show dynamic changes in glucose metabolism in response to checkerboard stimulation with a temporal resolution of 60sec. Subsequent studies have used this method to show task-*locked* FDG-*f*PET (time-locked to an external stimulus; Jamadar et al., 2019) and task-*related* FDG-*f*PET (not time-locked to an external stimulus; Hahn et al., 2016; 2018) metabolic activity. Using these methods, FDG-*f*PET temporal resolutions of 60sec have been obtained, which is a substantial improvement above bolus methods. Preliminary data from our lab show that the infusion method can provide temporal resolutions of between 20-60sec (Jamadar et al., 2018; current results)

Despite the promising results from the constant infusion method, the plasma radioactivity curves of these studies show that the infusion method is not sufficient to reach steady-state within the timeframe of a 90-min scan (Jamadar et al., 2019; Villien et al., 2014). In addition to the constant infusion procedure, Carson (2000) also proposed a bolus plus infusion procedure, where the goal is to quickly reach equilibrium at the beginning of the scan, and then sustain plasma radioactivity levels at equilibrium for the duration of the scan. Rischka et al. (2018) recently applied this technique using a 20% bolus plus 80% infusion; as expected the arterial input function quickly rose above baseline levels and was sustained at a higher rate for longer, compared to previous results using an infusion only procedure (e.g., Jamadar et al., 2019; Villien et al., 2014).

In this paper, we will describe in detail the acquisition protocol for acquiring high temporal resolution FDG-*f*PET scans using infusion only and bolus plus infusion radiotracer administration. Our protocols have been developed for use in a simultaneous MRI-PET environment with a 90-95min acquisition time (see Jamadar et al., 2019). While our group’s focus is the application of infusion methods for functional neuroimaging using BOLD-*f*MRI/FDG-*f*PET, these methods can be applied to any FDG-*f*PET study regardless of whether simultaneous MRI, BOLD-*f*MRI, computed tomography (CT) or other neuroimages are acquired.

## Protocol

This protocol has been reviewed and approved by the Monash University Human Research Ethics Committee, in accordance with the Australian National Statement on Ethical Conduct in Human Research (2007). Procedures were developed under the guidance of an accredited Medical Physicist, Nuclear Medicine Technologist (NMT) and clinical radiographer. Researchers should refer to their local experts and guidelines for the administration of ionising radiation in humans.

In the simultaneous MRI-PET environment, this procedure requires four personnel: a radiographer (RG) to run the scan, a nuclear medicine technologist (NMT) to oversee the administration of the radiotracer and acquisition of blood samples, a laboratory assistant (LA) to spin blood, and a research assistant (RA) responsible to oversee the experimental design and stimulus presentation. We use a commercial supplier for the production of the radiotracer. We report procedures for infusion only and bolus/infusion protocols. Figure 1 shows the flowchart of procedures in this protocol.

**Figure 1:**
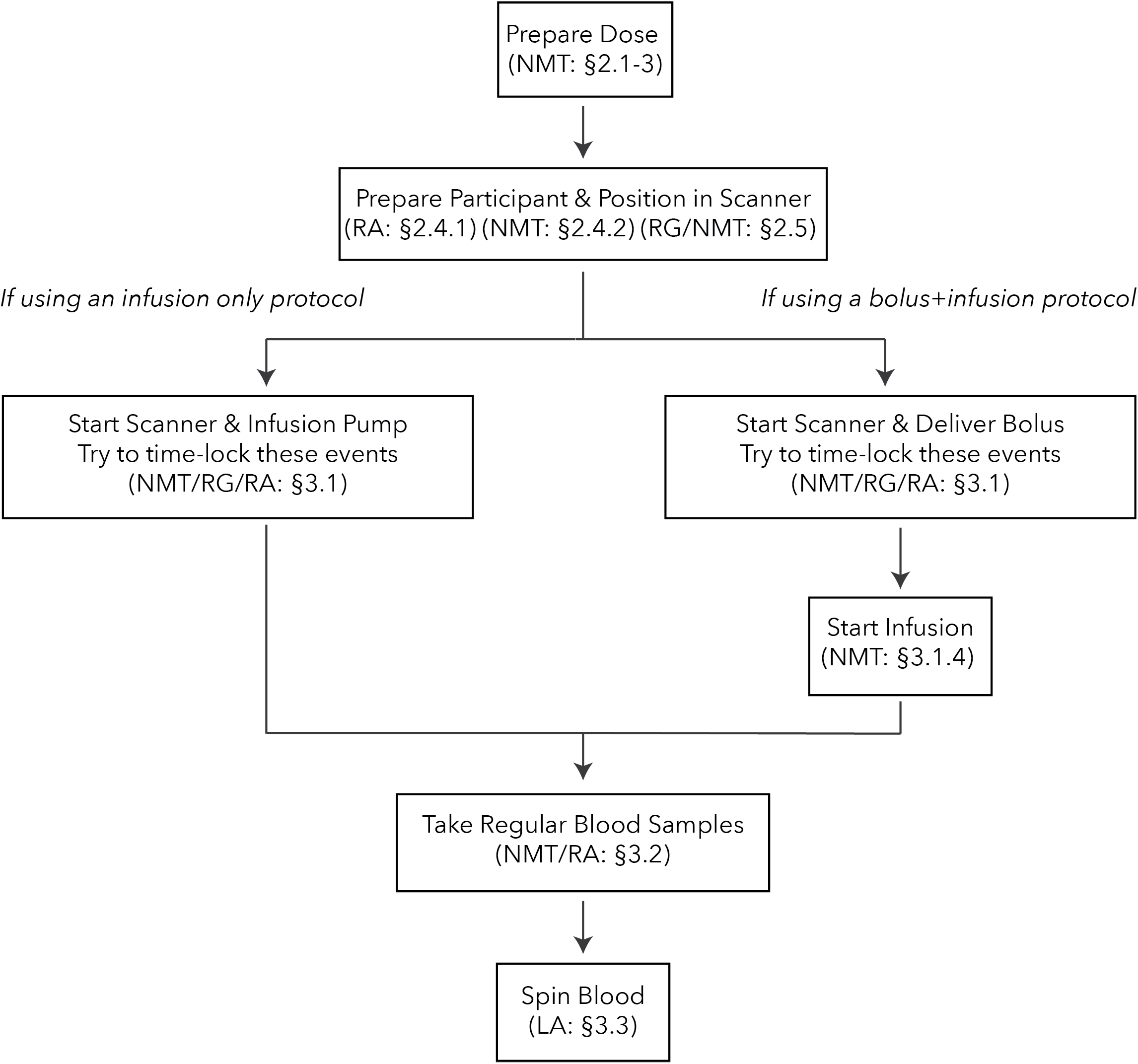
Flowchart of procedures for the infusion only (left) and bolus/infusion (right) protocols. In parentheses we list the staff member responsible for each procedure. Section identifiers refer to the sections in text where the procedure is described. Abbreviations: NMT: Nuclear Medicine Technologist; RA: Research Assistant; RG: Radiographer; LA: Lab Assistant.

### 1. Required equipment

Table 1 shows the Table of Materials for the scanner room, radiochemistry lab, and general materials.

**Table 1.** Table of Materials

| Where required | Supply | Preparation |
| --- | --- | --- |
| Scanner Room | 12-15 vacutainers | Remain in sterile packaging until required to put blood in tube |
|  | 12-15 10mL LH blood collecting tubes | Marked with the sample number (e.g., S1, S2...) and subsequently marked with the sample time (e.g., time 0 + x min [T<sub>0</sub>+x]) |
|  | 12-15 10mL pre-drawn 0.9% saline flushes | These are drawn up on the day of the study and capped with the ampoule that contained the saline |
|  | 12-15 5mL syringes | Remain in sterile packaging until ready to withdraw a blood sample |
|  | Gloves, waste bags, cello underpads ('blueys'), pen | All objects arranged on a plastic table/chair inside the scanner room on the same side as the arm from which the blood samples will be taken |
| Radiochemistry Lab | 1 18-gauge needle | Remain in sterile packaging until ready to inject <sup>18</sup>FFDG into the saline bag |
|  | 100mL 0.9% normal saline bag | Nil |
|  | ThermoScientific Heraeus Megafuge 16 centrifuge | Set to 2000RPM and 5mins. Acceleration and deceleration set to 8 |
|  | Well counter | Complete quality control and protocol set to 18F and 4mins. |
|  | Pipette | Set to 1000μL |
|  | 12-15 plasma counting tubes and pipette tips | Marked in the same manner as the LH blood tubes |
|  | 3 tube racks | Checked with a Geiger counter to ensure there is no radiation contamination on them |
|  | Gloves, waste bags, cello underpads ('blueys'), pen |  |
| Additional equipment required for the protocol | Synchronised clocks in scanner room, console and radiochemistry lab |  |
|  | Lead shields |  |
|  | Infusion pump and tubing | MR-compatible; e.g., BodyGuard 323 MR-compatible infusion pump (Caesarea Medical Electronics, Caesarea, Israel) |
|  | Lead bricks |  |
|  | Syringe shields |  |
|  | Geiger counter |  |

### 2. Preparation

(NB: Important points highlighted in yellow, as per Instructions to Authors)

2.1. Dose preparation (NMT)
  2.1.1. Calculate infusion volume that will be administered over the course of the scan. In our protocol, the rate of infusion is 0.01mL/second over 95 minutes. So, in a 95min scan, participants receive 0.01ml * 60sec * 95mins = 57mL.
  2.1.2. Calculate the dose that will be diluted into the administered solution (saline). In our protocol, we administer a total dose of 260MBq to the participant over 95mins. Decay correct 260MBq from the mid-infusion point (47.5mins) back to T0. Using equation 1, solve for A_0_

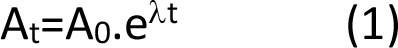 Where A_t_ is activity (MBq) at the mid-timepoint of the infusion; A_0_ is initial activity and *λ* is the radioactive decay constant specific to the tracer. For FDG, the value of *λ* is ≈ 0.693/T_1/2_. T_1/2_ is the half-life of ^18^F (110minutes). In our example, A_t_=260MBq; *λ* = 0.693/110; t=−47.5, so A_0_=350.942MBq.
  2.1.3. Calculate the required dose activity for the 100mL saline bag that will be used to administer the dose to the participant. For a 100mL saline bag, the dilution factor is volume of saline (100mL) divided by the infusion volume (57mL); i.e., 100mL/57mL = 1.754. So, the total dose required for the 100mL bag is A_0_*dilution factor; i.e., 350.942MBq*1.754 = 615.688MBq. Add this dose to the saline bag.
  2.1.4. Prepare the radiotracer.
    2.1.4.1. Prepare the priming dose. Withdraw 20mL from the bag into a syringe and cap it. Calibrate this 20mL syringe and affix label.
    2.1.4.2. Prepare the dose. Using a 50mL syringe, withdraw 60mL from the bag and cap with a red combi stopper. This syringe cannot be calibrated. Store both syringes in the radiochemistry lab until ready to scan.
    2.1.4.3. Prepare the reference dose. Fill a 500mL volumetric flask with approximately 480mL of distilled water. Draw up 10MBq of ^18^F FDG into a syringe, decay corrected to the scan start time (using equation 1) and add it to the flask. Top the volume up to the 500mL mark with more distilled water and mix thoroughly. Affix pre- and post-calibration labels of the syringe.
2.2. Prepare the scanner room (NMT)
  2.2.1. Once the participant is positioned in the scanner there is very little room to manipulate or salvage the line (for infusion or blood samples) if blockage occurs. The NMT should prepare the scanner room to minimise the chance of line blockage.
  2.2.2. The NMT should ensure all blood collecting equipment is within easy reach of the collection site. Cello underpads should be placed at the end of the cannula and on any surface that blood products and their containers will rest. Bins for regular waste and biohazardous waste should be within easy reach of the blood collection site.
2.3. Prepare the infusion pump (NMT)
  2.3.1. Set up infusion pump in scanner room on the side that will be connected to the participant. Build lead bricks around the base of the pump and place the lead shield in front of the pump.
  2.3.2. Connect the tubing for the infusion pump that delivers the infusion to the participant and ensure the correct infusion rate has been entered. For this protocol the rate is 0.01mL/second.
  2.3.3. The tubing needs to be primed before it is connected to the participant’s cannula. Connect the 20mL priming dose to the infusion pump. On the end of the tubing that will be connected to the participant, attach a 3-way tap and an empty 20mL syringe. Ensure the tap is positioned to allow the ^18^FFDG solution to flow from the priming dose through the tubing and collect into the empty syringe only.
  2.3.4. Pre-set the infusion pump to prime a volume of 15mL. Select the ‘prime’ button on the pump and follow the prompts to prime the line.
  2.3.5. Connect the 50mL dose syringe to the infusion pump in place of the priming dose. The 15mL primed dose on the 3-way tap can remain there until the participant is ready to be connected to the pump.
2.4. Participant preparation (NMT, RA, radiographer)
  2.4.1. The RA conducts consent procedures and the acquisition of additional measures (e.g., demographic surveys, cognitive batteries, etc.). NMT and radiographer conduct safety screens.
  2.4.2. Cannulate the participant.
    2.4.2.1. The most appropriate cannula varies across participants, but the ‘best’ vein should be reserve for blood collection; 22-gauge cannula is the preferred minimum size. Collect a 10mL baseline blood sample while cannulating. All saline flushes need to be disconnected under pressure to maintain patency of the line.
    2.4.2.2. Test participant blood sugar level and other baseline blood measures (e.g., haemoglobin) from the baseline sample.
2.5. Position the participant in the scanner (radiographer, NMT)
  2.5.1. The radiographer will position the participant in the scanner bore. For long scans, it is imperative to ensure participant comfort to reduce to risk of participant drop out, or motion artefact due to discomfort.
  2.5.2. The NMT must flush the cannula to ensure it is patent with minimal resistance before connecting the infusion line. Once connected the tubing can be lightly taped near the wrist. The participant must be instructed to keep their arm straightened. Supports such as foam or cushions are worthwhile for comfort. The NMT must also check the cannula that will be used for plasma samples to ensure that blood is able to be withdrawn with minimal resistance, and extension tubing to make the cannula more accessible whilst they are in the scanner is not leaking.
  2.5.3. Once positioned in the scanner bore, the NMT must check that they have suitable access to both cannulas.
  2.5.4. The NMT should notify the radiographer and RA if there are any issues with the blood collection cannula, infusion cannula or the infusion pump (e.g., occlusion, battery, extravasation) at any time during the scan.

### 3. Scan the participant

3.1. Start the scan (NMT, radiographer, RA)
  3.1.1. At the start of the scan, the NMT is positioned in the scanner room monitoring the infusion equipment. Ensure the NMT is wearing hearing protection.
  3.1.2. The radiographer performs the localiser scan to ensure that the participant is in the correct position. The details for the PET acquisition are checked (e.g. scan duration, Listmode data collection, correct isotope are the most important)
  3.1.3. The protocol is designed so that the PET acquisition will commence with the first MRI sequence. The radiographer prepares the first MRI sequence. The radiographer starts the MRI sequence; the start time of the 95-minute PET acquisition is time-locked to the start of the MRI sequence. If required, the NMT should deliver the bolus at the time of PET acquisition (Figure 1).
  3.1.4. Start the infusion pump: 30-seconds after the start of the PET acquisition, the radiographer signals to the NMT (e.g., via thumbs up sign) to start the pump. We start the infusion pump 30-seconds after scan start time to provide a safety buffer in case of scan failure. This also ensures that the first image taken during the PET scan indexes the brain prior to radiotracer administration for complete time activity curve data collection. The NMT will observe the pump to ensure it has started to infuse the 18F FDG and there is no immediate occlusion of the line.
  3.1.5. The RA is responsible for initiating any external stimulus at the agreed upon time (i.e., at the start of a functional run/experimental block).
  3.1.6. The RA is responsible for calculating the times for blood samples. An example Record Form is shown in the Supplement. The RA should calculate the predicted time of each blood sample, and provide copies of this to the NMT and lab assistant (LA). The RA is responsible for ensuring the NMT takes the blood samples at approximately the correct time, and ongoing monitoring equipment (e.g., infusion pump, stimulus) for any signs of errors.
3.2. Take regular blood samples (NMT, RA)
  3.2.1. If acquiring MRI scans simultaneously with PET scans, the NMT should wear hearing protection when entering the scanner room.
  3.2.2. The NMT must wear gloves and the tip of the cannula swabbed clean. While the cannula site dries, open a 5mL and a 10mL syringe, vacutainer and a 10mL saline flush.
  3.2.2. The NMT must wear gloves and the tip of the cannula swabbed clean. While the cannula site dries, open a 5mL and a 10mL syringe, vacutainer and a 10mL saline flush
  3.2.3. Using the 5mL syringe, withdraw 4-5mL of fresh blood and discard the syringe in the biohazard waste.
  3.2.4. Using the 10mL syringe, gently withdraw 10mL of blood. At the 5mL mark, provide an agreed-upon sign (e.g., thumbs up) to the RA. The RA will mark this time on the Record Form (Supplement) as the ‘actual’ time of sample.
  3.2.5. Connect the 10mL syringe to the vacutainer and then deposit the blood into the relevant blood tube.
  3.2.6. Quickly flush the cannula with 10mL of saline, disconnected under pressure, to minimise any chance of line clotting.
  3.2.7. The blood sample should be immediately taken to the radiochemistry lab for analysis.
3.3. Spin the blood (LA).
  3.3.1. The LA should have all equipment ready (Table 2) and be wearing gloves. It is preferable to have 3 racks set out for the samples: one for blood tubes (pre- and post-sample), another for pipetting the sample, and the last for filled pipetted samples (pre- and post-counting). Regularly change gloves throughout the procedure, especially when handling the counting tube. If the LA has any radioactive plasma contamination on their gloves, it can be transferred to the counting tube and spuriously increase the number of recorded counts of the sample.
  3.3.2. The blood sample can be placed in the centrifuge at any time. All samples must be spun at 2000RPM for 5-mins.
  3.3.3. Once the sample has been spun, place the tube in the pipetting rack. Remove the tube cap so as not to disturb sample separation. Place a labelled counting tube in the rack.
  3.3.4. Ensure the tip is securely fastened to the pipette. Have a tissue ready for any drips. Steadily pipette 1000μL from the blood tube and transfer to the counting tube.
  3.3.5. Place the counting tube into the well counter and count for 4-mins. The counting start time should be recorded on the Record Sheet (‘measurement time’) for every sample. At later time-points during the scan, the LA will be performing each step in rapid succession to avoid a backlog of samples.
  3.3.6. Dispose of any blood product waste in biohazard bags.

### 4. Study-Specific Methods

Here, we report study-specific details for the representative results; these details are not critical to the procedure and will vary across studies.

#### 4.1 Participants & Task Design

Participants (n=3) underwent a simultaneous BOLD-*f*MRI/FDG-*f*PET study. The full results of the study will be reported in Ward et al. (forthcoming). As this manuscript focuses on the PET acquisition protocol, MRI results are not reported here. Participants received 260MBq of [18F]-FDG over the course of a 95-min scan. Participant 1 received the full dose as a bolus at the start of the scan. Participant 2 received the dose in an infusion-only protocol, and participant 3 received the same dose with a hybrid 50% bolus+50% infusion. For both infusion-only and bolus/infusion protocols, infusion duration was 50-mins.

The task was presented in an embedded block design (Jamadar et al., 2018). We have previously shown that this design provides simultaneous contrast for task-evoked BOLD-*f*MRI and FDG-*f*PET data. Briefly, the task alternated between 640sec flashing checkerboard blocks and 320sec rest blocks. This slow alternation provides FDG-*f*PET contrast. Within the 640sec checkerboard blocks, checkerboard and rest periods alternated at a rate of 20sec on/20sec off. This fast alternation, which is suited to BOLD-*f*MRI, will hopefully be detectible with FDG-*f*PET with future analysis and reconstruction advances.

#### 4.2 Image Acquisition & Processing

PET data was acquired in list mode. The MRI and PET scans were acquired in the following order (details provided only for images relevant to the current manuscript): (i) T1-weighted 3D MPRAGE (TA=7.01mins, TR=1640ms, TE=2.34ms, flip angle=8°, FOV=256×256mm^2^, voxel size=1×1×1mm^3^, 176 slices; sagittal acquisition), (ii) T2-weighted FLAIR (TA=5.52mins), (iii) QSM (TA=6.86mins), (iv) gradient field map (TA=1.03mins), (v) MR attenuation correction Dixon (TA=0.39mins, TR=4.1ms, TE_in phase_=2.5ms, TE_out_ phase=1.3ms, flip angle=10°), (vi) T2*-weighted echo-planar images (EPIs) (TA=90.09mins), P-A phase correction (TA=0.36mins) and (vii) UTE (TA=1.96min). The onset of the PET acquisition was locked to the onset of the T2* EPIs.

T1-weighted structural images were neck-cropped using FSL-robustfov (Jenkinson, Beckmann, Behrens, Woolrich, & Smith, 2012), bias corrected using N4 (Tustison et al., 2010) and brain extracted using ANTs (Avants, Klein, Tustison, Woo, & Gee, 2010; Avants, Epstein, Grossman, & Gee, 2008). T1-weighted images were non-linearly normalised to MNI space using ANTs (Avants et al., 2011).

We examined dynamic FDG-fPET results with bin size 16-sec. All data were reconstructed offline using Siemens Syngo E11p and corrected for attenuation using pseudoCT (Burgos et al., 2014). Ordinary Poisson - Ordered Subset Expectation Maximization (OP-OSEM) algorithm with Point Spread Function (PSF) modelling (Panin, Kehren, Michel, & Casey, 2006) was used with 3 iterations, 21 subsets and 344×344×127 (voxels size: 2.09×2.09×2.03 mm^3^) reconstruction matrix size. A 5-mm 3-D Gaussian post-filtering was applied to the final reconstructed images.

Spatial realignment was performed on the dynamic FDG-fPET images using FSL MCFLIRT (Jenkinson, Bannister, Brady, & Smith, 2002). A mean FDG-PET image was derived from the entire dynamic timeseries and rigidly-normalised to the individual’s high-resolution T1-weighted image using ANTs (Avants et al., 2011). The dynamic FDG-fPET images were then normalised to MNI space using the rigid transform in combination with the non-linear T1 to MNI warp.

First-level models were estimated using SPM12 (Wellcome Centre for Human Neuroimaging) with the event time-course (checkerboard on, fixation) modelled as the effect of interest. Average uptake across the visual cortex (Brodmann areas, BAs, 17, 18, 19) was included as a covariate. The model did not include global normalisation, high-pass filter or masking threshold. An explicit mask of BAs 17, 18, 19 created in WFU-Pickatlas (ref; dilation 3) was included in the model to restrict the model estimation to regions of interest. T contrasts were used to estimate parameter maps of the individual-level activity, liberally thresholded at T=0.1 (uncorrected), k=50 voxels.

## Representative Results

### Plasma Radioactivity Concentration

The plasma radioactivity concentration curve for each participant is given in Figure 2.

**Figure 2:**
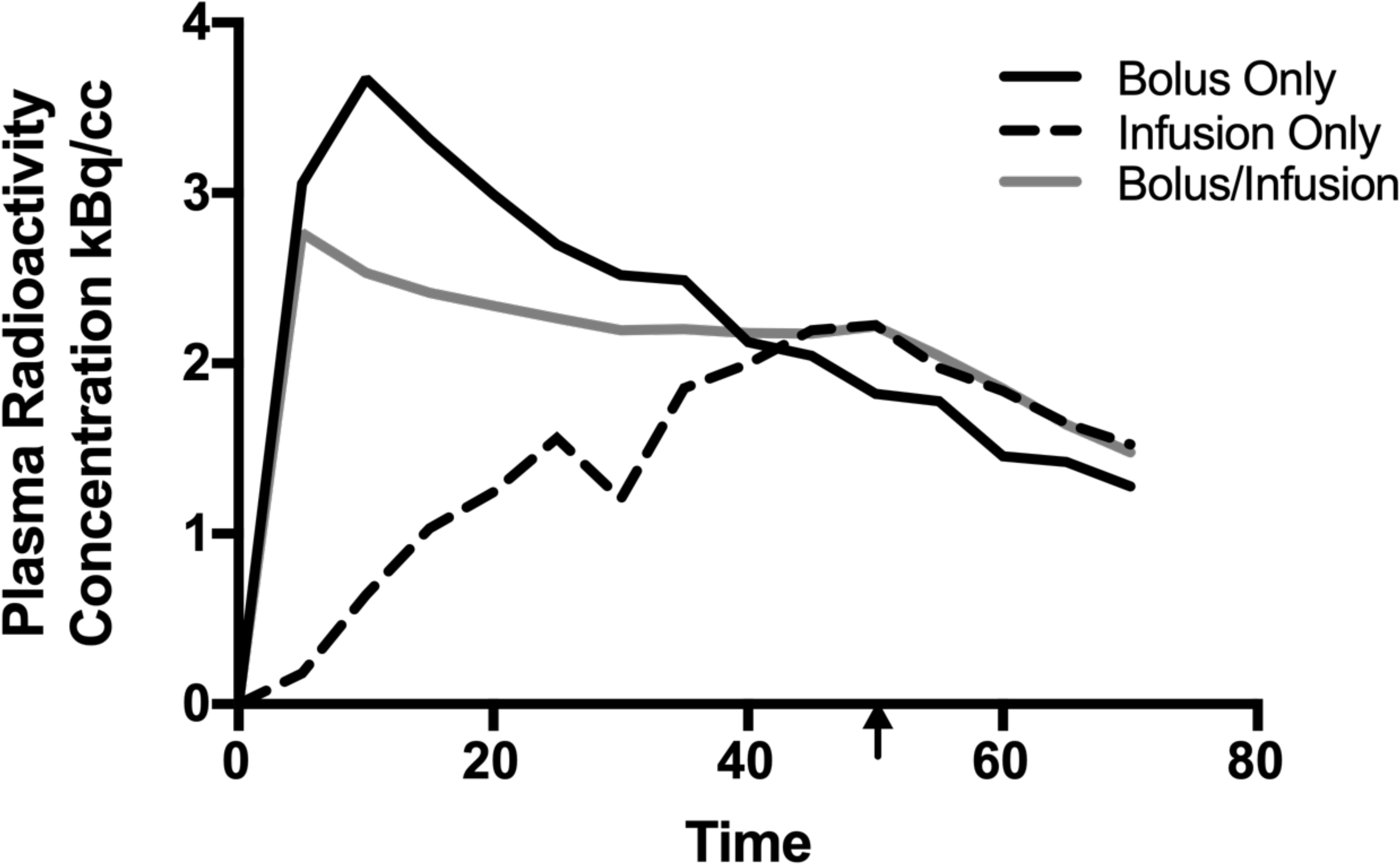
Plasma radioactivity curves for the three participants. Decay was corrected to the time the blood was sampled. The arrow indicates the cessation of the infusion for the infusion-only and bolus/infusion protocols.

The largest peak plasma radioactivity concentration (3.67kBq/cc) was obtained using the bolus method. The peak occurs within the first 5-mins of the protocol (note that the first blood sample was taken at 5-mins post-bolus), and the concentration quickly decreases. By the end of the recording period, the plasma radioactivity was 35% of the peak (1.28kBq/cc). The infusion only protocol reached maxima (2.22kBq/cc) at 50-mins, the end of the infusion period. By the end of the recording period, the concentration was sustained at 68% of its peak (1.52kBq/cc). Like the bolus only protocol, the bolus/infusion protocol reached its peak plasma radioactivity concentration (2.77kBq/cc) within the first 5-mins. By the end of the recording period, bolus/infusion concentration was at 53% of the peak (1.49kBq/cc).

Qualitatively, plasma radioactivity levels were sustained for the longest duration in the bolus/infusion protocol. Both infusion-only and bolus/infusion protocols show an apparent reduction in radioactivity when the infusion period ends (50-mins). Comparing bolus-only and bolus/infusion protocols, plasma radioactivity was smaller in bolus-only vs. bolus/infusion by 40-mins post-injection. Critically, the plasma radioactivity was minimally varied for a period of approximately 40 minutes in the bolus/infusion protocol. In contrast, neither the infusion-only or bolus-only protocol exhibit a sustained period of consistent activity.

### Image-Derived Uptake

Individual-level parameter maps, image-derived uptake and timecourses are shown in Figure 3.

**Figure 3:**
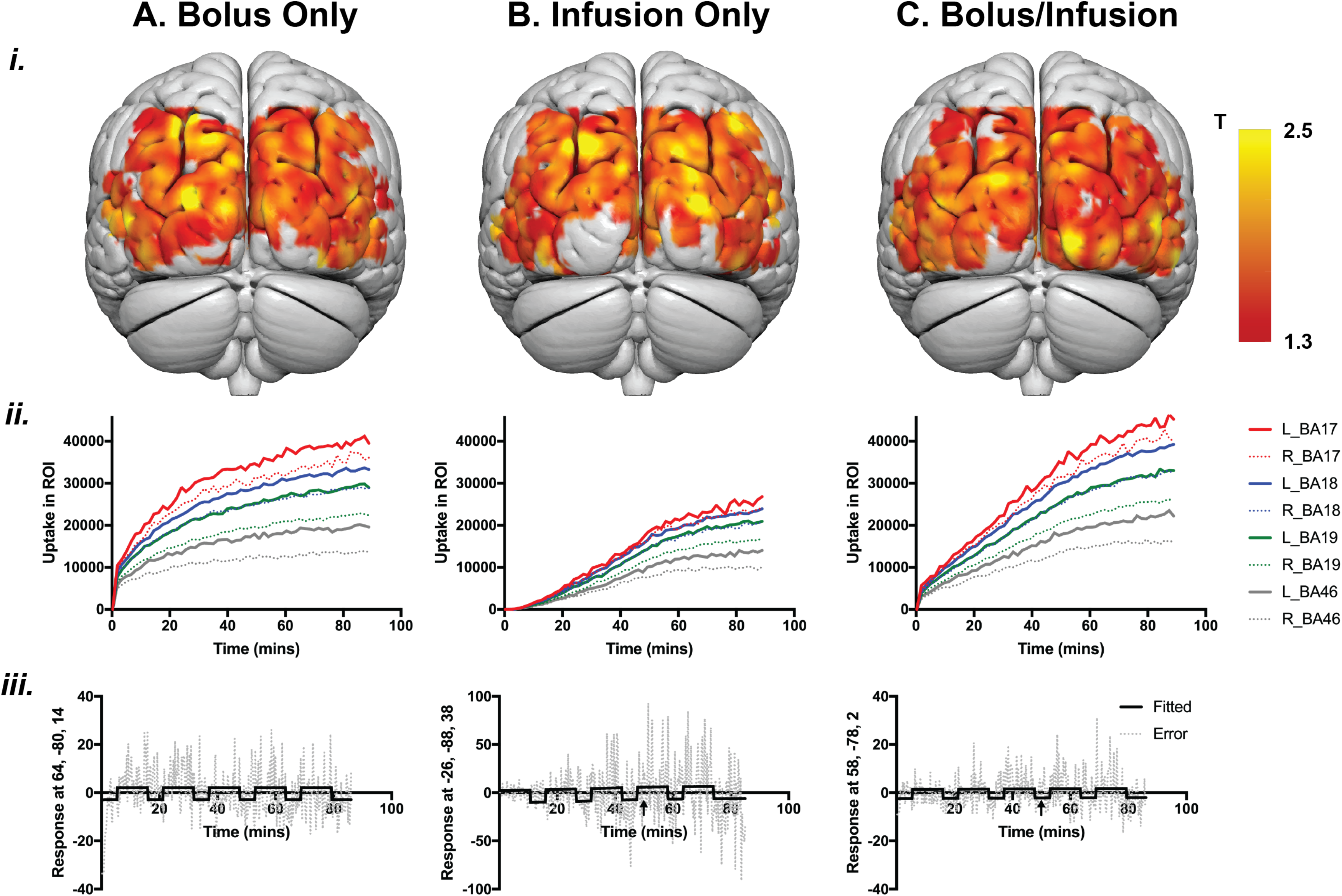
**(i)** Individual-level statistical parameter (T) maps for each of the three subjects, thresholded at p (uncorrected) < 0.1, k=50 voxels. **(ii)** Image-derived uptake across the visual cortex in four Brodmann areas (BAs): three occipital (BA17, 18, 19) and one frontal (BA46) control areas. **(iii)** Model fit and error across time for the peak of activity in each subject; arrow shows the end of the infusion period. For bolus-only (Aiii) peak activity MNI coordinate (64, −80, 14) T= 4.73; infusion-only (Biii) peak activity MNI coordinate (−26, −88, 38) T=4.14; bolus/infusion (Ciii) peak activity coordinate (58, −78, 2) T=5.05. Note that for the infusion-only protocol, the model could not be estimated for the first rest period due to very low signal. Also note the larger scale for the infusion-only protocol compared to the bolus-only and bolus/infusion protocols.

Figure 3ii shows the FDG uptake in bilateral visual cortex (BA17, 18, 19) and in a control region (dorsolateral prefrontal cortex, BA 46) for the three administration protocols. Across all protocols, the largest uptake is in primary visual cortex (BA17), followed by upper visual regions (BAs 18 and 19), as expected from the task design. The lowest uptake was in dorsolateral prefrontal cortex (BA46). Interestingly, for each ROI, uptake was larger in the left than the right hemisphere.

In the bolus protocol, there is a sharp increase in signal following the bolus. The slope of the uptake is relatively fast in the next 20-30 mins, but the rate of uptake reduces in the remainder of the measurement period. In the bolus/infusion protocol, there is a sharp increase in uptake at the start of the scan that is of smaller magnitude than in the bolus-only protocol, and the uptake continues at a comparatively faster rate for the duration of the scan. By the end of the recording period, the bolus/infusion protocol shows a larger uptake than the bolus only protocol, at least in the primary visual cortex (BA17). By comparison, the infusion-only protocol shows low signal for the first 40-mins of the scan, and the peak uptake is substantially lower than the bolus-only or bolus/infusion protocol. Uptake is fastest in the first ~50-mins of the scan, and slows for the remainder of the recording period.

### Parameter Maps and Timecourses

Figure 3i shows the individual-level T maps for the three administration protocols. Figure 3iii shows the timecourse at the peak voxel for each subject. Note that for the infusion only protocol (Figure 3Biii), the scale is larger than for the bolus only and bolus/infusion protocols. Furthermore, the model could not be estimated for the first rest period of the infusion only protocol, due to the very low signal; thus, the timecourse is shown from the start of the first checkerboard period.

To visualise the task effects across time, we extracted the timecourse data for each subject, and calculated the inverse of the coefficient of variation (mean/s.d.) for each block. The inverse of the coefficient of variation approximates signal-to-noise ratio. As can be seen from Figure 4, signal decreased roughly linearly across the recording period for the bolus-only protocol, increased roughly linearly for the infusion-only protocol, and was relatively flat for the bolus/infusion protocol.

**Figure 4:**
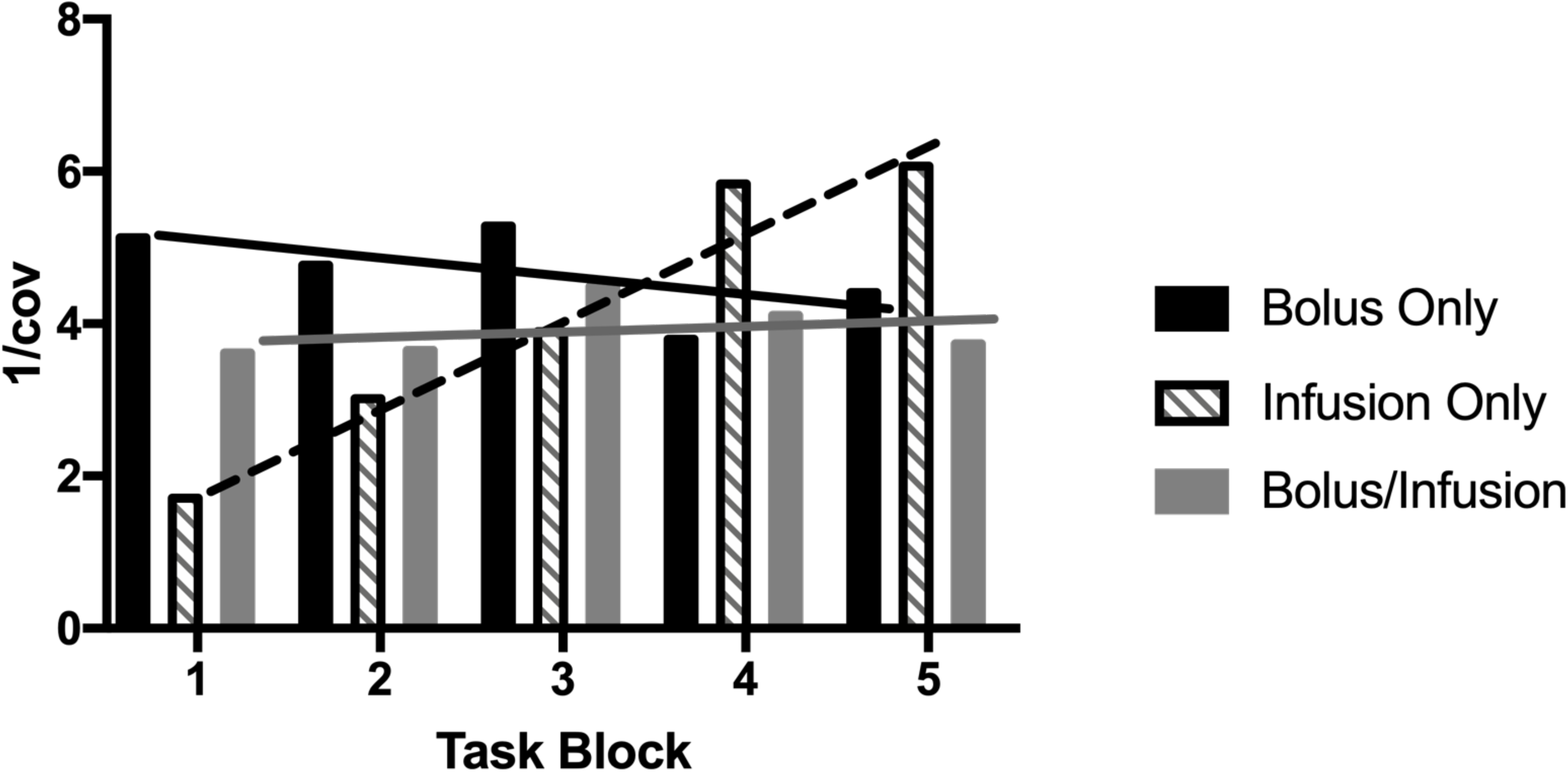
Signal to noise ratio across the recording period. Plot shows inverse of the coefficient of variation (mean/s.d.) of the first eigenvariate of the activity within the peak voxel in each checkerboard block.

## Discussion

FDG-PET is a powerful imaging technology that indexes cerebral glucose metabolism. To date, the vast majority of neuroscience studies using FDG-PET have used a traditional bolus administration approach, with a static image resolution that represents the integral of all metabolic activity over the course of the scan (Chen et al., 2018). In this manuscript, we have described two alternative radiotracer administration protocols: the infusion-only (e.g., Jamadar et al., 2019; Villien et al., 2014) and the hybrid bolus/infusion (e.g., Rischka et al., 2018) protocols. We have demonstrated a superior temporal resolution of 16-sec, time-locked to a stimulus, at the individual-level for the three protocols.

The critical point in the protocol is the start of the scanning protocol. At this point, the beginning of the PET acquisition must be time-locked to the beginning of the BOLD-fMRI sequence (if using simultaneous MR-PET), as well as the start of the stimulus presentation. Stimulus onsets and durations must be able to be locked to the onset of the scan for the first-level models. In the bolus-only protocol, the bolus should be delivered at the beginning of the PET acquisition to capture the peak signal (Figure 4). In the infusion-only protocol, the beginning of the infusion should be locked to the PET acquisition, to ensure accurate modelling of the uptake at the first-level. In the bolus/infusion protocol, the bolus should be time-locked to the PET acquisition, with the infusion starting at a known, short period, after the bolus. In order for the procedures to flow correctly within this short time period, each of the staff members (NMT, radiographer, research assistant) should be adequately prepared prior to the start of the scan (Figure 1). Dress-rehearsals are recommended to choreograph the timing of this critical stage.

To date, we have tested approximately 60 subjects using one of these protocols (the largest number using the infusion-only protocol). We have found that the likeliest cause of subject attrition or acquisition failure are due to:

A. being unable to cannulate the participant due to difficulty finding veins – to address this, we ask all participants to drink at least 2 glasses of water before the scan. If only one cannula can be achieved, we forego blood sampling for that participant.
B. Participants being unable to complete the scan. Unlike MRI, the PET acquisition cannot be paused then restarted. The most common causes of this are due to toilet breaks and difficulty with thermal regulation. Participants have reported that the requirement to consume water before the scan increases the need to urinate, thus all participants are required to use the bathroom prior to scanning. Participants have also reported that the infusion of the tracer leaves them feeling very cold – and the shivering reflex is triggered in some people. We address this by using a disposable quilt for all participants during the scan.

Results were shown at the individual subject level for the three administration protocols. As expected, the blood plasma radioactivity concentration (Figure 2) had the largest peak for the bolus-only protocol, but the most sustained radioactivity in the bolus/infusion protocol. The plasma concentration was lowest for the infusion-only protocol; and for both infusion-only and bolus/infusion, the concentration decreased at the time the infusion ceased. Image-derived uptake across the four bilateral ROIs (Figure 3ii) was consistent with the plasma-based measures, with largest uptake evident in the primary visual cortex, and smallest in the dorsolateral prefrontal cortex, as expected. Qualitatively, rate of uptake was largest for the longest period of time in the bolus/infusion protocol, and smallest in the infusion-only protocol. It is possible that the infusion-only protocol would yield better results in a longer experiment (>50mins) however this would likely increase the rate of participant attrition.

In the first-level models, model error was much greater in the infusion-only protocol compared to the bolus-only and bolus/infusion protocols (Figure 3iii). Signal-to-noise during the task periods (Figure 4) confirmed that the most sustained signal across the recording period was obtained using the bolus/infusion protocol.

In summary, we present alternative methods of FDG radiotracer administration for high-temporal resolution FDG-PET, with a resolution of 16sec. This temporal resolution is substantially improved over current standards in the literature (1-min: Hahn et al., 2016; 2018; Jamadar et al., 2019; Villien et al., 2014. Note Rischka et al., 2018 achieved stable results with a frame duration of 12sec using 20/80% bolus/infusion). The bolus/infusion protocol presented here appears to provide the most stable signal for the most sustained period of time, compared to the bolus-only and infusion-only protocols.

## Acknowledgements

Jamadar is supported by an Australian Council for Research (ARC) Discovery Early Career Researcher Award (DECRA DE150100406). Jamadar, Ward and Egan are supported by the ARC Centre of Excellence for Integrative Brain Function (CE114100007). Chen and Li are supported by funding from the Reignwood Cultural Foundation.

Jamadar, Ward, Carey & McIntyre designed the protocol. Carey, McIntyre, Sasan & Fallon collected the data. Jamadar, Ward, Parkes & Sasan analysed the data. Jamadar, Ward, Carey & McIntyre wrote the first draft of the manuscript. All authors have reviewed and approved the final version.

## Supplement

Example participant record form. In our protocol, the RA is responsible for recording the time of bolus & infusion start, then calculating the time of blood samples. The RA then provides copies of this form to the NMT and Lab Assistant. During the course of the experiment, the RA records the times that the samples were taken for subsequent decay correction. The Lab Assistant records the time of measurement and the measurement values in the Notes section.

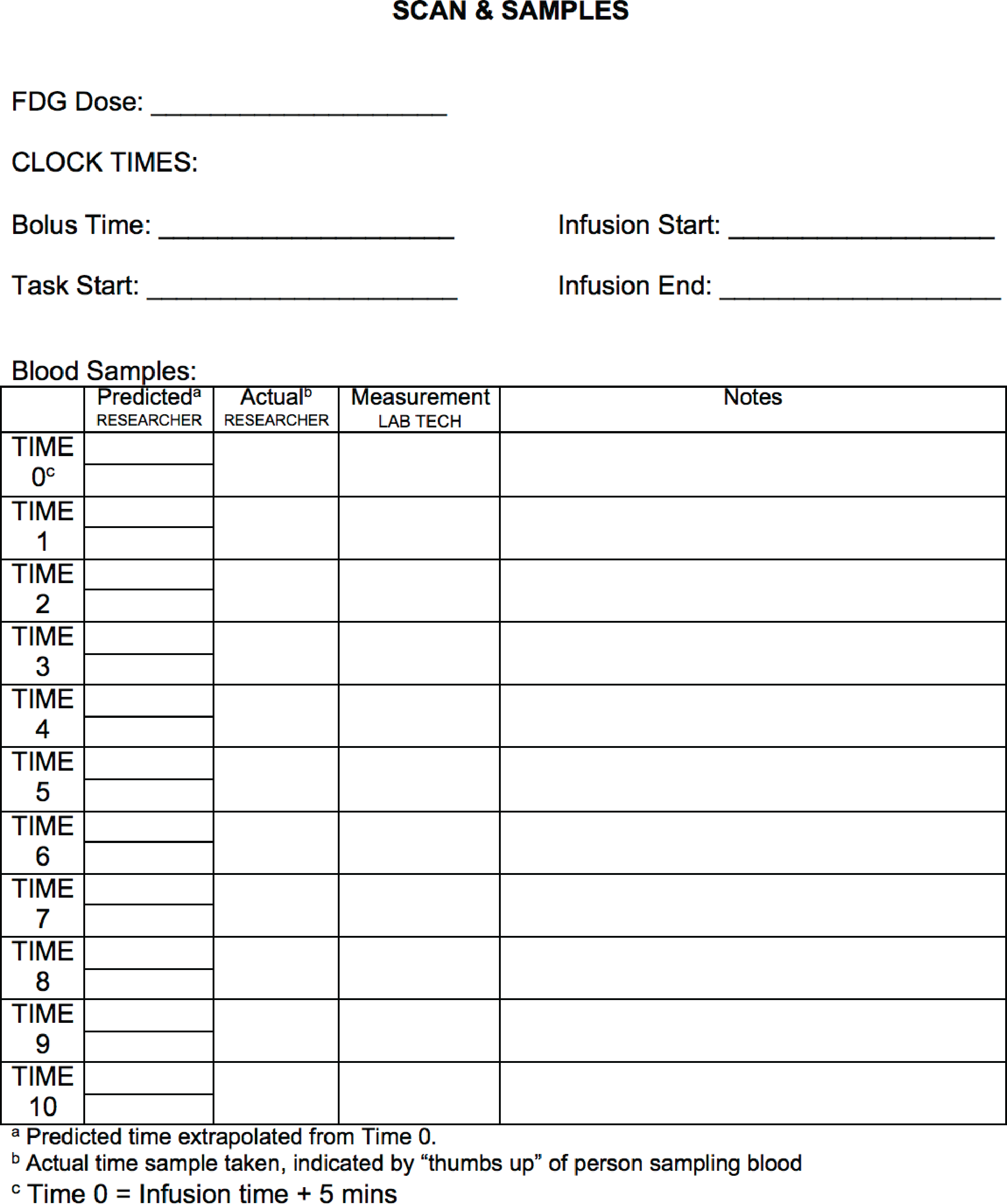

## Footnotes

1 It is beyond the scope of this manuscript to explore Simpson’s Paradox; but briefly, it refers to the phenomenon that brain-behaviour relationships calculated across-subjects are not necessarily indicative of the same relationships tested within-subjects. Thus, on the basis of between-subject correlations, one might infer that there is a positive brain-behaviour relationship, when in fact the relationship is negative when tested within-subjects. See Roberts et al. (2016) for a full exploration of this effect in neuroimaging.

